# Sucker recruitment patterns on the octopus arm during chemotactile exploration

**DOI:** 10.64898/2026.11.23.768135

**Authors:** Haleigh C. Anderson, Kendra C. Buresch, Raphaelle Tremblay, Ashley J. Gendreau, Max Lu, Aidan Sasser, Ava Agharabi, Corrinna Weyhrich, Roger T. Hanlon

**Author notes:** These authors contributed equally to this work (Haleigh C. Anderson, Kendra C. Buresch).

## Abstract

Octopuses possess highly flexible arms lined with chemo- and mechanosensory suckers that mediate exploration, prey capture, and object manipulation. How individual suckers communicate with adjacent and distant suckers to accomplish such tasks is poorly understood. We developed a behavioral protocol with *Octopus bimaculoides* to investigate inter-sucker recruitment once an individual sucker encountered an object of interest. We filmed the suckers on individual arms interacting with agarose disks embedded with prey extract and analyzed recruitment dynamics along the two longitudinal rows of suckers on each arm. We observed 18 recruitment patterns involving different sequences of ipsilateral and contralateral suckers.

Octopuses most frequently recruited ipsilateral neighboring suckers along the same row, yet many patterns included contralateral suckers in diverse combinations. Recruitment occurred more rapidly toward the arm tip than toward the arm base, and local coordination was ultrafast (mean 155.9 ms between sucker attachments). This directionally organized, ipsilaterally biased, yet flexible recruitment is consistent with distributed neural control along the octopus arm, which may allow octopuses to adapt sucker coordination to the demands of manipulating different items, and may inspire innovation in sucker arrangement and coordination in soft robotic arms.

**SUMMARY STATEMENT:** Octopus suckers are recruited ultrafast (1/2 second) in many combinations, but preferentially along the same row and toward the arm tip, offering insight into peripheral arm control and bioinspired soft robotics.

## INTRODUCTION

Octopuses are sophisticated cephalopods with eight exceptionally flexible neuromuscular appendages specialized for chemotactile exploration and manipulation (Wells, 1978; Hanlon and Messenger, 2018). Despite their excellent vision, which facilitates many behaviors, many shallow-water octopus species largely rely on chemotactile exploration during foraging. This “touch-taste” strategy (Wells, 1963) is enabled by dozens of suckers lining each arm that contain specialized chemo-and mechanoreceptors as well as local neural circuitry capable of processing sensory information and coordinating sucker activity (Fouke and Rhodes, 2020; Grasso, 2008; Graziadei and Gagne, 1976; Hanlon and Messenger, 2018; van Giesen et al., 2020; Olson et al., 2025a).

Octopuses have a complex, highly distributed nervous system, with the majority of neurons located within the arms rather than the central brain (Young, 1963; Young, 1971;Hochner, 2012; Richter et al., 2015; Gutnick et al., 2020). This allows for coordinated movement between arms through a nerve ring that connects to the axial nerve cord of each arm (Chang and Hale, 2023).

Each sucker contains a ganglion and neural connections that enable it to send and receive information, and sensory signals can propagate along the axial nerve cord and between suckers, allowing different parts of the arm to perform coordinated actions simultaneously (Grasso, 2008; Bagheri et al., 2020; Graziadei and Gagne, 1976; Hanlon and Messenger, 2018; Young, 1971; Bennice et al., 2025). Previous studies using amputated arms demonstrated that sucker recruitment can occur independently of the central nervous system, indicating that local peripheral circuits are sufficient for coordinating some aspects of sucker activity (Rowell, 1963; Bagheri et al., 2020). Coordinated sucker movements support important behaviors such as object manipulation and substrate locomotion (Altman, 1971; Mather, 1998; Nesher et al., 2020). Because these behaviors place different demands on the suckers, recruitment may need to be adaptable rather than stereotyped. Such coordination likely reflects neural connections between individual suckers (Neacsu and Crook, 2024), and recent studies suggest that the axial nerve cord may also coordinate actions between suckers (Olson and Ragsdale, 2023; Olson et al., 2025b; Olson and Ragsdale, 2026).

Despite this anatomical and functional evidence for local coordination, the dynamics and flexibility of inter-sucker recruitment in intact, freely behaving octopuses remain poorly quantified. Preliminary work in *O. bimaculoides* and *E. dofleini* reported that stimulating one sucker influences its neighbors and that responses vary with distance and position along the arm (Hadjisolomou and Grasso, 2010; Nair et al., 2013), and sucker recruitment has been proposed to shape arm search behavior (Sivitilli et al., 2023). However, it remains unknown (i) whether sucker recruitment is directionally organized along the arm in freely behaving animals, (ii) if recruitment differs between ipsilateral and contralateral rows, (iii) how recruitment speed differs between sequences traveling toward the arm tip and those traveling toward the base, and (iv) the degree to which octopuses can recruit suckers in multiple combinations or instead rely on a stereotyped pattern.

We developed a behavioral assay with *Octopus bimaculoides* to quantify sucker recruitment dynamics *in vivo*. Animals inserted individual arms into an opaque chamber to interact with agarose disks embedded with prey extract, allowing close-up filming and visualization of sucker recruitment. From these videos we quantified: 1) the frequency of ipsilateral and contralateral recruitments progressing toward the arm tip and toward the body using sequences of 5 sequential sucker attachments, 2) the speed of these successive sucker recruitments, and 3) the diversity of sucker recruitment sequence patterns. A follow-up experiment was designed to examine variation in these sucker dynamics across different regions of the arm (proximal, medial, and distal).

Together, these metrics provide insight into how sensory information may propagate between suckers, reveal functional organization within the octopus arm, and indicate whether sucker coordination is flexible or follows a set pattern.

## RESULTS

In our initial experiment (Experiment 1), octopuses were observed sweeping their arms around the inside of the box until they encountered the agarose disks. Octopuses used the distal portion of their arm 80% of the time when interacting with the agarose disks. The disks were initially contacted with an individual sucker (or suckers) and additional suckers were recruited as octopuses wrapped their arm around the disk. Sucker recruitment was most often sequential (recruitment of an adjacent sucker 66% of the time), but in some cases the sucker adjacent to the last recruitment was skipped and the next sucker along the row was recruited instead (34% of the time). There was no difference in the direction of the first recruitment – whether it progressed toward the arm tip or toward the body (number of recruitments toward arm tip = 27.5, number of recruitments toward body = 22.5; X^2^_9,100_ = 10.80, p > 0.05).

In our follow-up experiment (Experiment 2), designed to compare sucker dynamics in different arm regions, octopuses quickly swam to the corner where the agarose disks were positioned and began interacting with them after being introduced to the tank. Octopuses swept their arms across the corner tank walls, wrapping the distal portion of the arm around the disks while also engaging the medial and proximal regions. In some cases, multiple regions of the same arm wrapped around a single disk, while in others a second arm was recruited. They frequently interacted with disks on both sides of the corner simultaneously. Suckers on the distal part of the arm were recruited most frequently (58.5% of all observations), but suckers on the medial (38.3% of all observations) and proximal (3.2% of all observations) parts of the arm were also observed to be recruited. The distal region of the arm was most often observed wrapping around the rim of the disk, while the proximal region often attached to the front face of the disk. Though proximal wrapping did occasionally occur, it generally involved fewer than five suckers, therefore data from this region was excluded from our analyses.

### Ipsilateral vs Contralateral Sucker Recruitments

#### Distal arm region

There were no differences in the frequency of recruitments in the distal arm region between Experiments 1 & 2 (toward arm tip: U = 5376, p = 0.8590; toward body: U = 4048, p = 0.4195), so data were pooled for comparison of frequency of ipsilateral vs contralateral recruitments. Ipsilateral sucker recruitments occurred more frequently than contralateral sucker recruitments in the distal region of the arm – for recruitments occurring both toward the arm tip and toward the body. Sequences toward the arm tip had an average of 3.95 ipsilateral and 1.06 contralateral recruitments per 5-sucker-recruitment sequence (n = 417, U = 2165, p < 0.0001; Fig. 1A). Sequences toward the body had an average of 3.25 ipsilateral and 1.75 contralateral recruitments per sequence (n = 374, U = 8913, p < 0.0001; Fig. 1A).

**Figure 1.**
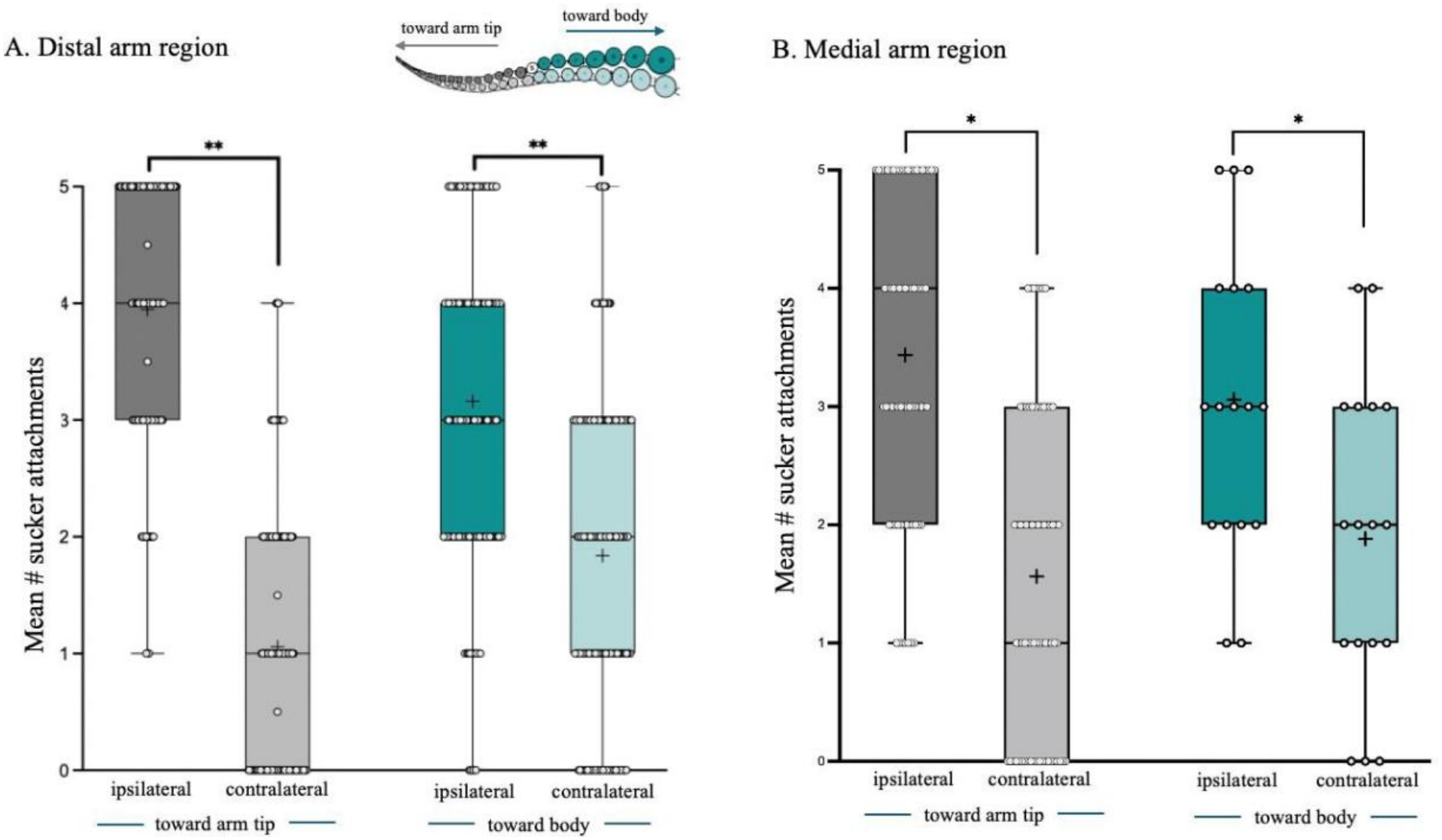
Sucker recruitments along the ipsilateral row occurred more frequently than sucker recruitments to the contralateral row - both toward the arm tip and toward the body in the distal and medial arm regions. **(A)** Distal arm region: Sequences toward the arm tip had an average of 3.95 ipsilateral and 1.06 contralateral recruitments per sequence (n = 417, U = 2165, p < 0.001). Sequences toward the body had an average of 3.25 ipsilateral and 1.75 contralateral recruitments per sequence (n = 374, U = 8913, p < 0.001). **(B)** Medial arm region: Sequences toward the arm tip had a mean of 3.44 ipsilateral and 1.56 contralateral recruitments per sequence (n = 142, U = 882, p < 0.001) and toward the body had an average of 3.06 ipsilateral and 1.88 contralateral recruitments (n = 34, U = 79, p = 0.022). Error bars represent the standard error of the mean, open circles represent individual data points, and asterisks indicate significant differences.

#### Medial arm region

In the medial region of the arm ipsilateral sucker recruitments also occurred more frequently than contralateral recruitments both toward the arm tip and toward the body.

Sequences toward the arm tip had an average of 3.44 ipsilateral and 1.56 contralateral recruitments per sequence (n = 142, U = 882, p <0.001; Fig. 1B). Sequences toward the body had an average of 3.06 ipsilateral and 1.88 contralateral recruitments per sequence (n = 34, U = 79, p = 0.022; Fig. 1B).

### Sucker Recruitment Sequence Patterns

Octopuses interacted with the agarose disks using 18 different sucker recruitment patterns that used both ipsilateral and contralateral recruitments. Three sucker recruitment sequence patterns were most common (both toward the arm tip and toward the body; Fig. 2A). In Experiment 1, the most frequently observed pattern was a pattern in which all five sucker recruitments in the sequence were ipsilateral relative to the previously attached sucker (toward the arm tip = 39%, n = 93, X^2^_31,93_ = 574.03, p < 0.001; toward the body 29%, n = 97, X^2^_31,97_ = 194.92, p < 0.001; Sequence 1, Fig. 2A). The ipsilateral only pattern was also the most frequently observed pattern overall in Experiment 2 (occurring 31% of the time overall). The next most frequently observed sequence was a pattern in which the first 4 sucker recruitments were ipsilateral and the 5th recruitment was contralateral (toward the arm tip = 19%, toward the body = 17%; Sequence 2, Fig. 2A). The third most frequent sequence had a pattern in which the first 3 recruitments were ipsilateral, the 4th recruitment was contralateral, and the final recruitment was ipsilateral relative to the 4th recruitment (toward the arm tip = 15%, toward the body 8%; Sequence 3, Fig. 2A). Sequences 2 and 3 occurred with similar frequency (toward arm tip Sequence 2 vs. Sequence 3: X^2^_1,93_ = 1.0, p > 0.05; toward body Sequence 2 vs. Sequence 3: X^2^_1,97_ = 0.059, p>0.05). Overall, 15 additional patterns were observed that involved different combinations of ipsilateral and contralateral recruitments; all were observed less than 5% of the time (Fig. 2B)

**Figure 2.**
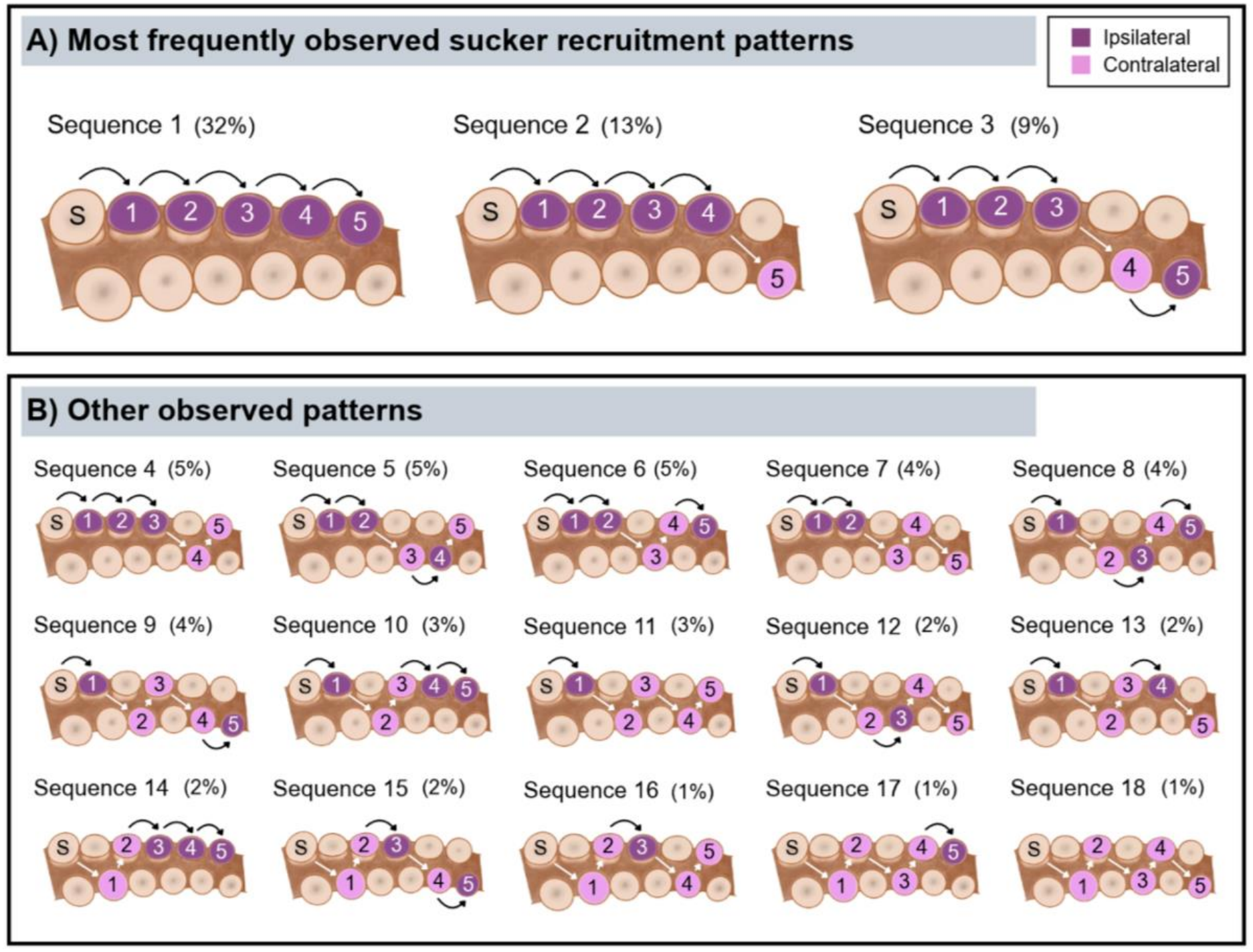
Observed diversity of sucker recruitment patterns. Octopuses interacted with the agarose disks using 18 different sucker recruitment patterns. **(A)** Most frequently observed patterns: Sequence 1 was the most frequently observed pattern (toward the arm tip = 32% of recruitments); all sucker recruitments in this sequence were ipsilateral relative to the previously attached sucker. In Sequence 2 the first 4 recruitments were ipsilateral and the 5^th^ was contralateral (13% of recruitments). In Sequence 3 the first 3 recruitments were ipsilateral, the 4^th^ recruitment was contralateral, and the final recruitment was ipsilateral relative to the 4^th^ recruitment (9% of recruitments). **(B)** Other observed sucker recruitment patterns: 15 additional patterns were observed; percent occurrence of each pattern is listed above the illustration of each pattern.

### Speed of Sucker Recruitment

#### Distal arm region

There was no difference in the speed between ipsilateral and contralateral sucker recruitments (mean time between ipsilateral recruitments = 110 ms, mean time between contralateral recruitments = 112 ms; p > 0.05) for the distal arm region, so data were pooled to determine if the direction of recruitment (toward the arm tip vs. toward the body) affected recruitment speed. Recruitment toward the arm tip occurred faster than recruitment toward the body for Experiment 1 (toward the arm tip = 91.62 ms, toward the body = 135.2 ms; n = 1000, U = 108175, p = 0.0002; Fig. 3A) and Experiment 2 (toward the arm tip = 128.3 ms, toward the body = 201.3 ms; n = 980, U = 95612, p < 0.0001; Fig. 3A).

**Figure 3.**
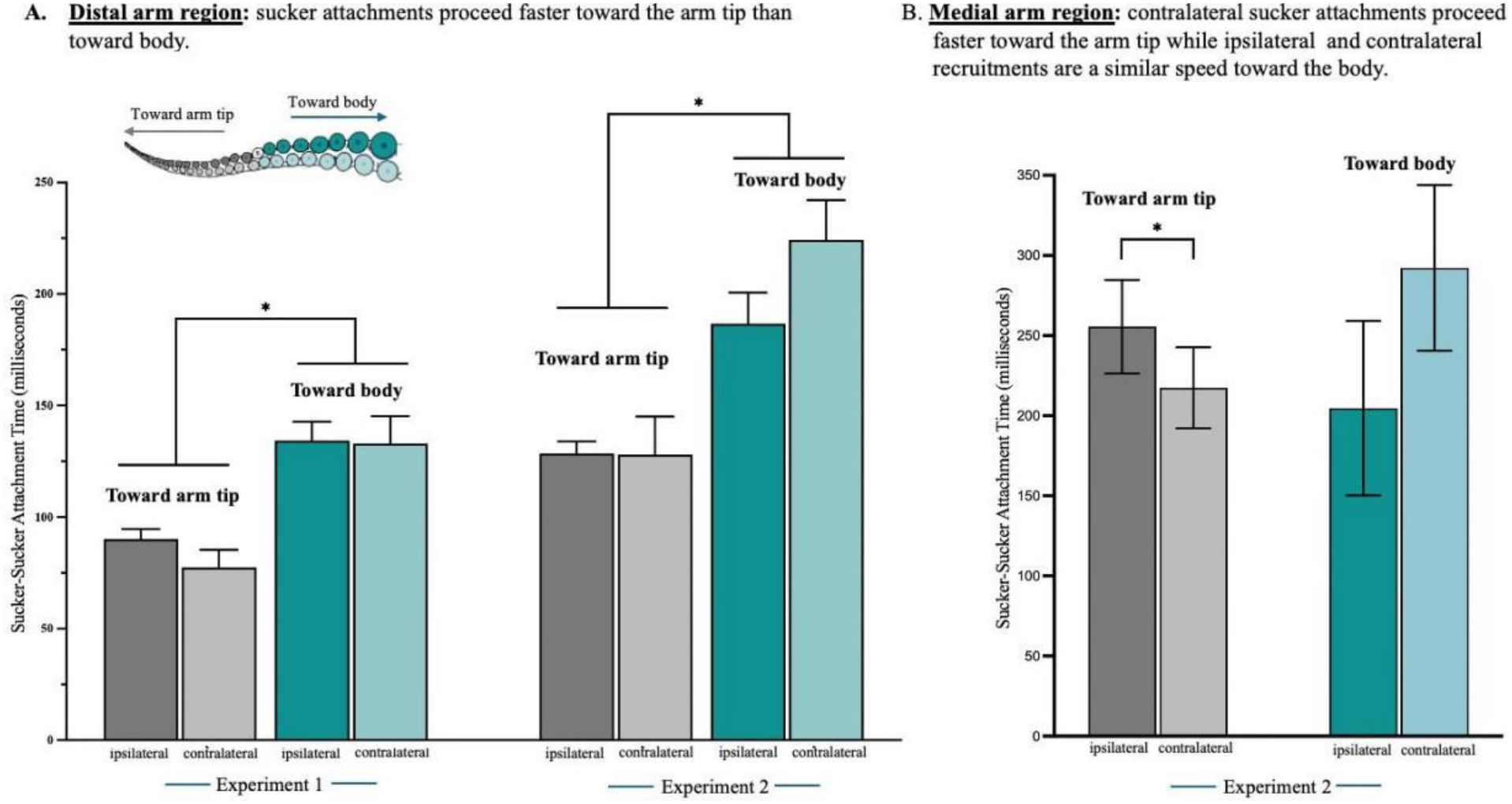
Speed between sucker recruitments in the distal and medial arm regions. **(A)** Distal arm region: Time between sequential sucker recruitments was faster toward the arm tip than towards the body for both Experiments 1 & 2 (Exp 1: U = 108175, p = 0.0002; Experiment 2: U = 95612, p < 0.0001). **(B)** Medial arm region: Contralateral recruitments were faster toward the arm tip (U = 4632, p = 0.0011), while there was no difference in the timing between ipsilateral and contralateral sucker recruitments toward the body (U = 1451, p = 0.1156). Error bars represent the standard error of the mean and asterisks indicate significant differences.

#### Medial arm region

The medial region of the arm showed more complex recruitment speed patterns than the distal region of the arm. For sequences toward the arm tip contralateral sucker recruitments occurred faster than ipsilateral recruitments; however, for recruitments occurring towards the body there was no significant difference in the timing between ipsilateral sucker recruitments and contralateral recruitments, but this could be due to a low sample size yielding low statistical power (toward arm tip mean ipsilateral = 255.5 ms, mean contralateral l = 217.4 ms; U = 4632, p = 0.0011; toward body mean ipsilateral = 204.7 ms, mean contralateral = 292.2 ms; U = 1451, p = 0.1156; Fig. 3B).

## DISCUSSION

This study demonstrates that sucker recruitment along octopus arms is ipsilaterally biased and most frequently propagates faster toward the distal tip than toward the base of the arm. Three recurring recruitment patterns featured predominantly ipsilateral sequences, and the most common was a fully ipsilateral sequence. This suggests that coordination between suckers follows consistent spatial biases, favoring communication within the same row and facilitating more rapid activation toward the arm tip. The predominance of ipsilateral sequences supports the idea that adjacent suckers are linked by local neural circuits that enable efficient coordination. However, the full set of 18 observed patterns, which included contralateral recruitment, indicates that this bias is not fixed and that octopuses can flexibly recruit suckers in other combinations.

While the precise neural pathways underlying sucker coordination remain unresolved (Olson and Ragsdale, 2023), our findings are consistent with neighboring suckers along the same row being functionally linked through local circuitry that facilitates short-range coordination. Although ultrastructural reconstructions have revealed prominent contralateral connectives linking sucker rows (Neacsu and Crook, 2024), these connectives may not be the primary route for the local recruitment that predominated here and instead may enable the less frequent contralateral recruitment patterns we observed. Ipsilateral pathways, potentially finer in scale and less readily resolved in EM reconstructions, may mediate the communication between adjacent suckers that characterized most recruitment sequences. Recent molecular and morphological evidence that sucker ganglia contain both sensory and motor neurons (Olson et al., 2025a) further supports the potential for such localized inter-sucker coordination. Together, these findings are consistent with a multi-layered organization in which short-range ipsilateral connections promote local recruitment, while contralateral connections support integration across sucker rows (Kuuspalu et al., 2022; Olson and Ragsdale, 2026). This organization could also account for the flexibility we observed: a default ipsilateral bias, with contralateral connectivity providing a substrate for the less frequent patterns that recruited suckers across rows. Such an arrangement may allow octopuses to adjust sucker coordination to the demands of manipulating different items.

In addition to preferential recruitment within the same row, sucker recruitment speed differed depending on position and direction along the arm. In general, recruitments propagated more quickly toward the arm tip, particularly within the distal region of the arm. This regional pattern is consistent with earlier reports that suckers proximal and distal to a stimulated sucker respond differently (Nair et al., 2013), although those experiments stimulated a single sucker rather than an intact arm interacting with a prey taste. This distal acceleration may reflect structural and neurochemical heterogeneity along the axial nerve cord (ANC) (Winters-Bostwick et al., 2024). Neuronal segments of the ANC decrease in size toward the distal tip while maintaining one segment per sucker (Olson et al., 2025b), potentially facilitating faster communication among more closely spaced distal modules. Regional variation in sucker diameter, which decreases distally, may also reduce conduction distance between adjacent suckers (Olson et al., 2025b). The medial region of the arm showed somewhat different recruitment speed patterns than the distal region, consistent with these underlying anatomical and neurochemical gradients. These structural gradients along the arm may therefore shape not only where recruitment occurs, but how quickly coordinated activation spreads.

While sensory and motor neurons are intermingled in the ANC (Gutfreund et al., 2006; Olson et al., 2025b), the faster distal recruitment seen here may reflect a difference in how motor commands versus sensory signals travel along the arm. Although sensory signals have been shown to propagate more rapidly toward the body (Zullo et al., 2011; Chang and Hale, 2023), the faster distal sucker recruitment observed here may instead reflect coordinated motor activation among neighboring suckers. Accelerated sucker recruitment toward the arm tip may be functionally advantageous given that this region is primarily responsible for prey capture and the initiation of the “fetching” motion that delivers food to the mouth (Sumbre et al., 2006; Richter et al., 2015). Rapid distal sucker recruitment could, therefore, reflect specialization of the arm tip for efficient prey acquisition and manipulation.

In addition to neural organization, the biomechanics of sucker recruitment also likely influence recruitment patterns. Sucker adhesion strength and type vary with behavioral context (Grasso, 2008; Röckner et al., 2023), and sucker attachment and release follow stereotyped, strategy-dependent patterns (Röckner et al., 2023; Röckner, 2026). The sucker recruitment patterns used for chemotactile foraging, such as when octopuses explore crevices and rocks in search of prey (Ambrose and Nelson, 1983; Mather and O’Dor, 1991; Forsythe and Hanlon, 1997; Leite et al., 2009; Mather et al., 2014; Hanlon and Messenger, 2018), may therefore differ when octopuses rely on visual cues or interact with objects of different shapes. Future work examining sucker recruitment across varied sensory and mechanical contexts will help determine the generality of these directional and ipsilateral biases.

This study provides the first analysis of the position, directionality, and speed of sucker recruitment in octopus arms in freely-behaving octopuses and revealed a clear bias toward adjacent ipsilateral recruitment and faster propagation toward the distal tip of the arm. These findings suggest neuroanatomical linkages between neighboring suckers that allow for rapid, directionally tuned coordination. Rather than acting as independent units, suckers appear to participate in organized recruitment patterns that likely enhance efficient prey capture and object manipulation. The additional recruitment combinations we observed alongside this dominant pattern suggest that this coordination is flexible and may allow octopuses to adapt sucker recruitment to the demands of manipulating different items. Understanding this local coordination provides insight into how complex, adaptive behavior emerges from distributed neural systems in flexible appendages. Beyond their biological significance, these results could inform the design of octopus-inspired soft robotic systems, where understanding how adhesion and control are distributed along flexible arms remains a central challenge (Mazzolai et al., 2012; Laschi et al., 2016; Yue et al., 2025; Del Dottore et al., 2026).

## MATERIALS AND METHODS

### Experimental Subjects

Ten *Octopus bimaculoides* (67 – 296.5 g) were collected from the Southern California coast and acclimated to laboratory conditions at the Marine Biological Laboratory in Woods Hole, MA. Octopuses were housed individually in 80-l aquaria supplied with a constant flow of filtered ambient seawater, bubble aeration, and maintained on a 12-hour light/dark cycle. Water temperature was held at 15 ± 0.5 °C and pH at 8.1 ± 0.1. Tanks were enriched with ceramic mug dens, artificial plants, and children’s toys. Prior to use as experimental subjects, octopuses were fed daily on a varied diet (*Carcinus maenas, Hemigrapsus sanguineus, Mytilus edulis)*.

### Experiment 1: Box Design

Octopuses (n = 10) were trained to interact with agarose disks embedded with prey extract (a mixture of shrimp and crab) presented inside a custom-built plexiglass box. One side of the box had a 3 cm or 1.5 cm circular opening, to allow access to the interior. The 3 cm opening was used exclusively during training, while the 1.5 cm opening was used during experimental trials to isolate individual arms for detailed behavioral recording. Agarose disks were affixed to a glass wall located opposite the circular opening, which was positioned flush against the side of the aquarium. This design enabled unobstructed video capture of sucker activity during interaction with the disks. To visually occlude the interior of the box and prevent the camera from focusing on the animal’s body, all box walls, excluding the glass wall, were covered with white felt in all experimental trials. The box was secured to the corner of the aquarium using three magnets.

### Experiment 1: Box Training Protocol

During training, octopuses were presented with a prey item (*Hemigrapsus sanguineus*) inside the plexiglass box and were considered successful if they removed the prey within the allotted time (Table 1). Training occurred twice daily, and progression to the next stage required three consecutive successful trials. The first three stages of training were conducted in the animal’s home tank. In the first stage, octopuses were given a clear-sided plexiglass box with a 3 cm opening and a live crab affixed to the wall opposite the opening. No time limit was imposed. In the second stage, the box remained the same, but a time limit of two hours was imposed. In the third stage, the time limit was reduced to one hour. The fourth stage occurred in a 75-l experimental tank using a whited-out box with a 1.5 cm opening and a one-hour time limit.

**Table 1.** Experiment 1 training and experimental protocol.

| Stage | Opening Size | Time Limit | Number of Trials | Item | Location |
|---|---|---|---|---|---|
| **1** | 3 cm transparent | unlimited | 3 | Crab | Home tank |
| **2** | 3 cm white-out | 2 hr | 3 | Crab | Home tank |
| **3** | 3 cm white-out | 1hr | 3 | Crab | Home tank |
| **4** | 1.5 cm white-out | 1hr | 3 | Crab | Test tank |
| **Experiment** | 1.5 cm white-out | 45 m | Until 10 video clips are collected | Crab/Agarose | Test tank |

Following three consecutive successful trials in this stage, the octopus was considered fully trained. If the octopus failed three consecutive trials at any stage, it was returned to the previous stage. The number of days required to complete training varied by individual.

### Experiment 1: Experimental Protocol

Once fully trained, each octopus (n = 10) was presented with two boxes every other day in a 75-l experimental tank (Fig. 4A). One box was introduced in the morning and the other in the afternoon. Each day, one box contained a live crab, used for feeding and to incentivize continued interaction with the box, while the other contained six agarose disks (1.5 cm diameter and 0.5 cm thick) embedded with prey extract and affixed to the interior glass wall of the box. The order of box presentation (crab or agarose) was randomized, and each box remained in the tank for 45 minutes. Filming only occurred during agarose trials. To minimize external visual stimuli and reduce human influence, black curtains were placed around the tank during recording. Two rectangular Godox™ WL8P LED lights were placed underneath the corner of the tank to increase illumination inside of the box. Video recordings were captured at 4K resolution and 60 FPS using an iPhone 15 running Filmic Pro (Evernote Corporation, version 7). Each octopus completed ten experimental trials.

**Figure 4.**
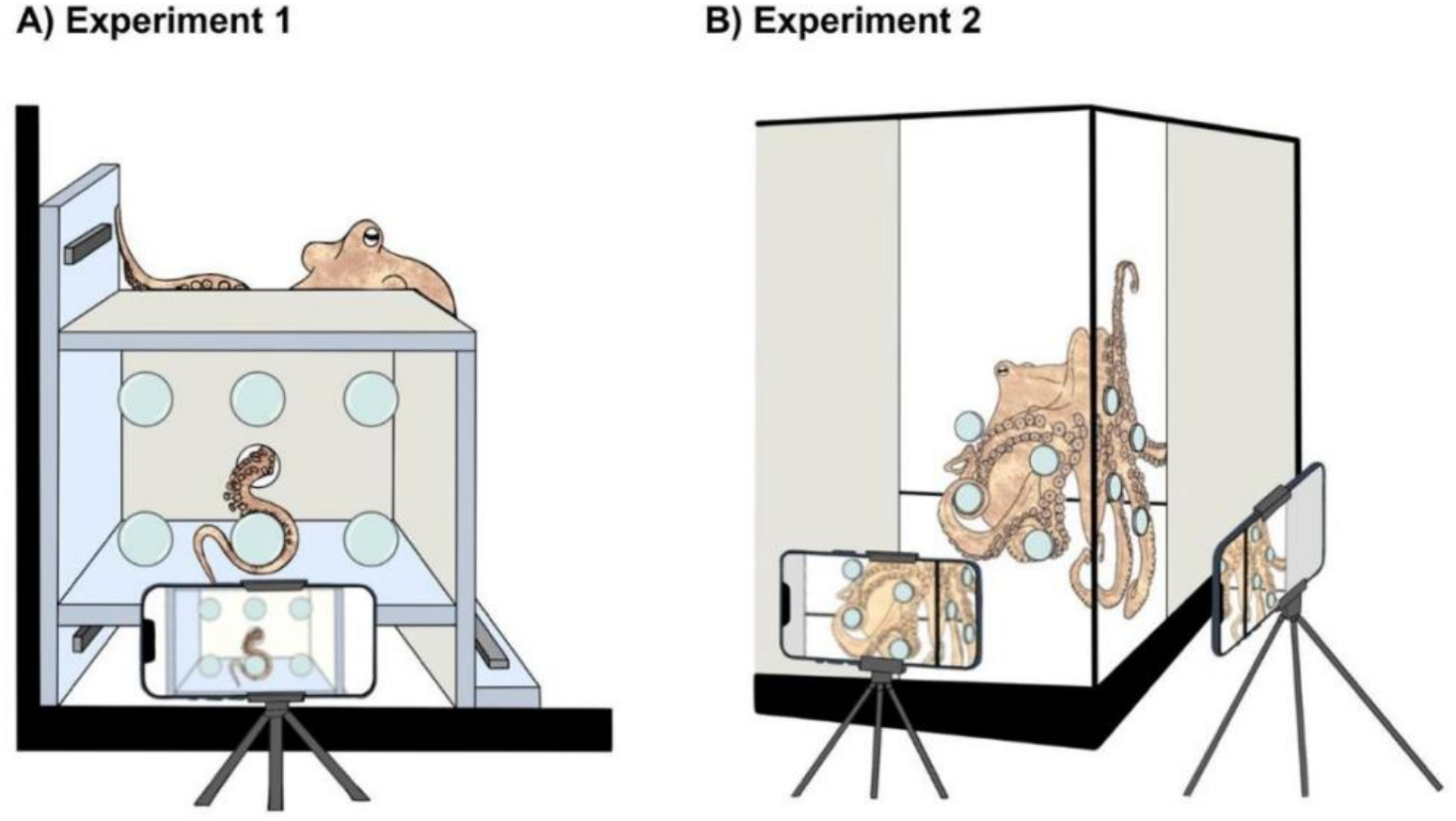
Experimental tank setup. **(A)** Experiment 1: A 75-l aquarium illuminated by two LED lights from beneath the tank; one camera is positioned for video recording. The octopus reaches through a hole in an opaque acrylic box to contact six agarose disks embedded with prey extract affixed to the inner wall of the box opposite the hole. **(B)** Experiment 2: a 75-l aquarium illuminated by two LED lights, one below and one above. Eight agarose disks embedded with prey extract are affixed to two adjoining walls (four per wall). Two cameras record the tank, one per wall, while the octopus freely explores and interacts with the disks.

### Experiment 2: Experimental Design

Since octopuses interacted with the agarose disks using only the distal region of the arm in 80% of video recordings in Experiment 1, we designed a second experimental protocol to enable direct comparison of sucker dynamics across the distal, medial, and proximal regions of the arm. This experiment was conducted using 5 individuals (137 – 317 g), and octopuses were allowed to freely explore the experimental tank and interact with the agarose disks using all 3 arm regions.

A 75-l glass aquarium filled with 15°C, filtered seawater served as the experimental tank. Eight agarose disks embedded with prey extract were affixed to two adjoining walls at one corner of the tank, with four disks per wall (Fig. 4B). To minimize external visual stimuli and reduce human influence, all sides of the aquarium were covered with opaque material, except for the two adjacent wall areas where the agarose disks were placed. The tank was illuminated from above with one Godox™ WL8P LED light and from below with two additional WL8P units to optimize visibility for video recording.

The octopus was introduced into the tank and allowed to freely explore for up to fifteen minutes. Two cameras (iPhone 14 and 15 Pro, using the Filmic Pro app; 4K resolution, 60 FPS) were mounted on tripods to record both walls. Two experimenters observed the animal’s behavior and activated the cameras once the octopus approached the agarose disks. Filming ceased after the octopus disengaged from the disks or removed all disks from the tank walls. If all disks were removed, the octopus was rewarded with a live crab (*Hemigrapsus sanguineus*) inside the tank. Each octopus (n = 5) completed ten experimental trials, conducted once per day, three times per week.

### Video Analyses

Video footage from both experiments was screened to identify short clips (1 – 2 seconds) in which the octopus contacted an agarose disk and both rows of suckers were clearly visible. Selected clips were analyzed frame-by-frame independently by two observers using Procreate (Savage Interactive, version 5.3.13). Suckers were scored as they attached to the agarose disk and were considered attached when the entire rim made contact. The frame number of each recruitment event was recorded. Observer scores were then compared to ensure accurate sucker identification and minimize observer bias. To score the sucker recruitment in each clip, a focal sucker (the first to attach within a given arm region) was identified. The first five sucker recruitments from the focal sucker were recorded in two directions: toward the arm tip and toward the body. Each recruitment was classified as ipsilateral or contralateral relative to the previously recruited sucker, allowing for 32 possible recruitment patterns.

In Experiment 1 ten video clips were selected for analysis from each octopus (n = 10), resulting in a total of 100 clips. Selected clips each contained five recruitments in two directions: toward the arm tip and toward the body (ten total recruitments), yielding two sequences per clip for a total of 200 sequences. Sequences progressing toward the arm tip and toward the body were analyzed separately to determine: 1) the frequency of ipsilateral versus contralateral sucker recruitment, 2) the latency between sucker recruitments and 3) patterns of sucker recruitment.

For Experiment 2 all video clips that had at least five sucker recruitments in either direction (toward the arm tip or toward the body) from each octopus (n = 5) were analyzed. Each clip was categorized according to the region of the arm used (distal, medial, or proximal). In the distal arm region, 109 sequences toward the arm tip and 87 sequences toward the body were analyzed. In the medial arm region, 71 sequences toward the arm tip and 17 sequences toward the body were analyzed. Only five sequences toward the arm tip were available from the proximal arm region, and no sequences toward the body met the inclusion criteria, so the proximal arm region was excluded from our analyses.

### Statistical Analyses

All statistical analyses were performed using GraphPad Prism 10 (GraphPad Prism version 10.4.1 for Mac, GraphPad Software, Boston, Massachusetts USA, www.graphpad.com). Recruitment pattern frequencies were analyzed using Pearson’s chi-squared tests. Pairwise comparisons between patterns were conducted using z-tests. Mann-Whitney U tests were used for comparisons between Experiments 1 and 2, to compare the frequency of ipsilateral versus contralateral sucker recruitments, and to compare sucker recruitment latencies.

## ACKNOWLEDGEMENTS

The authors would like to thank members of the Hanlon laboratory for their help with animal husbandry and data collection, with special thanks to Lee Gutman. We would also like to thank the members of the Marine Resources Department at the Marine Biological Laboratory for their support with water quality and seawater system maintenance. Thanks to Frank Grasso for insightful conversations about the design of sucker experiments.

## DATA AVAILABILITY

All data will be made available upon reasonable request to the corresponding author.

## DECLARATIONS

None

## ETHICAL APPROVAL

Cephalopods are not included in U.S federal regulations that govern the use of animals in research laboratories. However, the Marine Biological Laboratory regulates cephalopod research through their Institutional Animal Care and Use Committee (IACUC). The care of animals in this study adhered to the MBL’s institutional Cephalopod Care policy. This study was conducted under MBL IACUC protocol number 25-11K.

